# Genome-wide analyses reveal a strong association between LEPR gene variants and body fat reserves in ewes

**DOI:** 10.1101/2021.11.27.470213

**Authors:** Tiphaine Macé, Eliel González-García, Didier Foulquié, Fabien Carrière, Julien Pradel, Christian Durand, Sebastien Douls, Sara Parisot, Dominique Hazard

**Author notes:** Corresponding author: GENPHYSE, INRAE, 24 chemin de Borde Rouge, CS 52627, F-31326 Castanet-Tolosan, France. These authors contributed equally to this work (Tiphaine Macé, Dominique Hazard).

## Abstract

Among the adaptive capacities of animals, the management of energetic body reserves (BR) through the BR mobilization and accretion processes (BR dynamics, BRD) has become an increasingly valuable attribute for livestock sustainability, allowing animals to cope with more variable environments. BRD has previously been reported to be heritable in ruminants. In the present study, we conducted genome-wide studies (GWAS) in sheep to determine genetic variants associated with BRD. BR levels and BR changes over time were obtained through body condition score measurements at eight physiological stages throughout each productive cycle in Romane ewes (n=1034) and were used as phenotypes for GWAS. After quality controls and imputation, 48,513 single nucleotide polymorphisms (SNP) were included in the GWAS. Among the QTLs identified, a major QTL associated with BR levels during pregnancy and lactation was identified on chromosome 1. In this region, several significant SNPs mapped to the leptin receptor gene (LEPR), among which one SNP mapped to the coding sequence. The point mutation induces the p.P1019S substitution in the cytoplasmic domain, close to tyrosine phosphorylation sites. The frequency of the SNP associated with increased BR levels was 32%, and the LEPR genotype explained up to 5% of the variance of the trait. These results provide strong evidence for involvement of LEPR in the regulation of BRD in sheep and highlight it as a major candidate for improving adaptive capacities.

## Introduction

Breeding farm animals for adaptive traits is of growing interest for the improvement of livestock sustainability. Due to climate change and its associated challenges, it is expected that feed supply fluctuations will increase, both in terms of quantity and quality (Dumont *et al*. 2014). To cope with such nutritional challenges, animals rely on their energetic body reserves (BR) present in adipose tissues. Alternation of BR use and accretion periods, referred to as body reserve dynamics (BRD; i.e., lipid mobilization and accretion processes) in the present study, provides animals with a metabolic plasticity that allows them to respond to energetic challenges (for a review, see (Friggens *et al*. 2017)). Inclusion of BRD in future genetic programs is of particular interest to improve the adaptive capacities of animals and to optimize feeding management, especially for ruminants whose farming systems increasingly rely on rangeland and roughage resources (Phocas *et al*. 2016).

Some long-term energetic challenges are predictable and result in anticipatory changes in body reserves, e.g., the high energetic cost for lactation in mammals (for a review see (Friggens *et al*. 2017)). In this context, the temporal pattern of changes in BR can be genetically-driven. Genetic variability for adiposity has been previously described for several animal species, including humans (for a review, see (Speakman *et al*. 2008; Stachowiak *et al*. 2016; Abdalla *et al*. 2018; Bouchard 2021)). In ruminant species, total body energy content is a heritable trait in dairy cows throughout lactation (Banos *et al*. 2005). Heritability of BR levels estimated through the body condition score (BCS), a common proxy used to estimate BR in livestock and highly correlated with total body fat content, ranged between 0.08 and 0.45 for dairy cows and sheep (Russel *et al*. 1969; Mendizabal *et al*. 2010; Kenyon *et al*. 2014), depending on the breed and the physiological stage of the measurement (Koenen *et al*. 2001; Walkom *et al*. 2014a; Macé *et al*. 2018). Heritability of BR changes ranged from 0.01 to 0.16 in ruminants (Pryce *et al*. 2001; Dechow *et al*. 2002; Walkom *et al*. 2014b; Macé *et al*. 2018). These results indicated that BR levels and BRD were heritable traits in ruminants. Such genetic variability in temporal patterns of changes in BR could be used in genetic selection to improve adaptation and resilience. Recent developments in animal breeding provide the opportunity to include favorable polymorphisms for traits of interest through genomic selection. However, to our knowledge, while some studies have reported QTLs in sheep for fatness in the carcasses of lambs (Walling *et al*. 2004; Johnson *et al*. 2005; Matika *et al*. 2016; Garza Hernandez *et al*. 2018), no such studies have been undertaken for BR phenotypes on live productive females. Given the moderate to high heritability for BR phenotypes in sheep, we hypothesized the existence of some QTL regions underlying BR levels and BRD in productive ewes, among the 26 sheep chromosomes. Therefore, the objective of this study was to detect genetic variants associated with BR levels and BRD traits in ewes. These results should provide new insights to enhance our knowledge of fatness in mammals and molecular data that can be used to improve adaptive capacities in sheep by genomic selection.

## Materials and Methods

### Animals and management

The experiments described here fully comply with applicable legislation on research involving animal subjects in accordance with the European Union Council directive (2010/63/UE). The investigators who carried out the experiments were certified by the relevant French governmental authority as well as the INRAE La Fage Experimental Farm (agreement number A-12-203-1). All experimental procedures were performed according to the guidelines for the care and use of experimental animals and approved by the French Ministry of Education and Scientific Research and the local ethics committee CEEA-115 (Science and Animal Health) (approval number APAFIS#4597-2016031819254696.V3).

The experimental animals were Romane ewes reared at the INRAE *La Fage* Experimental Farm (*Causse du Larzac*, Saint-Jean Saint-Paul, southern France) between 2005 and 2019 (n=1034) (Ricordeau *et al*. 1992). Ewes were reared exclusively outdoors on approximately 280 ha of rangelands, in a flock of 250 reproductive females present each year. The main management features of this farming system have been previously described in detail (Molénat *et al*. 2005; Gonzalez-Garcia *et al*. 2014; Gonzalez-Garcia and Hazard 2016). Briefly, the farming system was based on a productive flock reared exclusively in extensive harsh conditions while limiting supplementation, in order to investigate the capacity of ewes to fend for themselves. In the autumn, before mating began, dry ewes successively grazed native and fertilized rangelands (6% of the total surface). A single mating period took place at the end of the autumn and first mating occurred at 8 or 20 months of age depending on the ewe’s live weight and experiment. During the winter and the second half of pregnancy, ewes were gradually supplemented with conserved feedstuffs (i.e., hay and silage produced on the farm) and barley due to the absence of grazeable biomass on the rangelands (for the detailed composition of the diet, see (Gonzalez-Garcia *et al*. 2014)). Lambing took place outdoors in the spring (April) and ewes suckled lambs for approximately 80 ± 4 days while they successively grazed fertilized and native rangelands. During the summer, dry ewes grazed the senescent vegetation due to drought on large paddocks containing a high proportion of shrubs (up to 30%). The Romane ewes produced an average of 2.2 live lambs per lambing in our conditions.

### Measurements

All the ewes were individually monitored for their body condition score (BCS), body weight (BW), subcutaneous back fat (BF) and muscle (BM) depth, and their pedigree information was recorded. All the data were recorded in an INRAE experimental database for sheep and goats: GEEDOC (https://germinal.toulouse.inra.fr/~mcbatut/GEEDOC/). BCS measurements were performed according the original scale described by Russel et al. (Russel *et al*. 1969) (i.e., ranging from 1, emaciated, to 5, obese) and the scale was adapted with a subdivision of 0.1 increments instead of 0.25. The same two operators systematically recorded the BCS measurements over the 15-year period and underwent regular training sessions for calibration. Subcutaneous back fat and muscle depths were measured by a realtime ultrasound system on the 12^th^ rib. The measurements used in this study were collected on a regular basis during one to three productive cycles, according to the following physiological stage schedule: mating (M, 15 days before mating), early pregnancy (Pa, 39 ± 11 days after mating), two-thirds of pregnancy (Pb, 101 ± 11 days after mating), lambing (L), early suckling (Sa, 17 ± 10 days after lambing), middle of the suckling period (Sb, 42 ± 10 days after lambing), weaning (-W, 80 ± 10 days after lambing) and post-weaning period (Wp, 149 ± 11 days after lambing). To characterize BRD, differences in BCS between pairs of physiological stages were calculated and analyzed (BCS-Pa:L, BCS-L:Sa, BCS-Pa:W, BCS-M:Pa, BCS-W:Wp and BCS-W:M; described by Macé et al. (Macé *et al*. 2019)). In addition to BCS, BW was also measured at all the stages described above, and BF and BM were measured at mating, lambing and weaning.

### Descriptive Statistics

Analyses of variance were carried out, taking the repeated measurements into account, using the MIXED procedure of the Statistical Analysis System (SAS version 9.4; SAS Institute Inc., Cary, NC, USA) to test relevant effects and interactions affecting phenotypes. The age at first lambing, the parity of the ewe, the litter size and the year of measurement were identified as fixed effects. The age at first lambing effect took account of ewes that lambed for the first time at 1 or 2 years of age (classes 1 and 2, respectively), the parity effect took account of first, second, third and more lambing (classes 1, 2 and 3, respectively), the litter size effect took account of the number of lambs born and suckled (class 0, empty ewes or lambing but without suckling lambs; class 1, ewes lambing and suckling singleton from L to W; class 2 ewes lambing more than singleton and suckling one lamb; class 3, ewes lambing and suckling twins; and class 4, ewes lambing and suckling more than twins). The first-order interactions between age at first lambing × litter size and parity × litter size were tested. An effect was considered significant if *P* < 0.05.

### Adjusted phenotypes

Phenotypes were adjusted for significant fixed effects using REML software with the following linear mixed model:

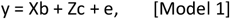

where y is the vector of observations for one of the BR traits; b is the vector of fixed effects consisting of age at first lambing, parity of the ewe, litter size, and year of measurement; c is the vector of random animal effects; and e is the vector of random residuals with incidence matrices X and Z. c and e were assumed to be normally distributed with means equal to zero and (co)variances Iσ^2^c, Iσ^2^e, respectively. I are identity matrices of appropriate size. Model 1 was fitted with an identity matrix I for random animal effects fitted instead with the pedigree-based relationship matrix, as performed in Mucha et al. (Mucha *et al*. 2018). The resulting estimated animal values were used as adjusted phenotypes for subsequent genomic analyses instead of estimating breeding values. This was done to take repeated measurements for each individual into account, considering animal effect as a permanent environmental effect, and to remove any influence of pedigree relationships (contributions of parents and relatives) on an animal’s value, as was previously observed with estimated breeding values (Ekine *et al*. 2014).

### SNP genotypes and quality control

Sheep were commercially genotyped with Illumina Ovine SNP15K (i.e., 16,560 Single Nucleotide Polymorphisms (SNP); low density (LD)), SNP50K (i.e., 54,241 SNPs, medium density (MD)) or SNP600K (i.e., 606,006 SNPs; high density (HD)) beadchips. Genotypes were established as part of the research projects: “SheepSNPQTL”, “COMPAGNE”, “RomaneIteDomum”, “iSAGE” and “SMARTER”. A total of 1,034 phenotyped female animals were genotyped and distributed as follow: 820 ewes were genotyped with the MD beadchip, 167 ewes were genotyped with the LD beadchip, and 47 ewes were genotyped with the HD beadchip. Among the phenotyped and genotyped females, 554 have their dam genotyped (i.e., 389 genotyped dams out of a total of 700 dams) and 965 have their sire genotyped (i.e., 49 genotyped sires out of a total of 60 sires). Dams and sires were genotyped with either MD or HD chips (374 parents and 64 parents, respectively). Concerning ewes genotyped with LD chip, both parents were genotyped.

Individuals with a call rate below 0.95 and with Mendelian inconsistencies were discarded (i.e., five genotyped and phenotyped ewes were removed). The SNPs were removed from further analyses if they were not in Hardy-Weinberg equilibrium, had a minor allele frequency below 1% or had a call rate below 0.98. PLINK software was used to detect incompatible genotypes between sires, dams and offspring (Chang *et al*. 2015). When the total number of incompatible SNPs was more than 2% of all SNPs, ewes were kept in the analyses and the parents in error were replaced by missing values (14 ewes concerned). For ewes genotyped with the HD chip, only SNPs present on the MD chip and LD chip were kept. The ewes genotyped with the LD chip were imputed to the MD chip using Fimpute software, as were LD chip SNPs absent in the MD chip (Sargolzaei *et al*. 2014). This resulted in a data set, used for QTL analyses, containing 1,034 ewes with BR phenotypes genotyped for 48,593 autosomal SNPs.

### QTL detection method

A genome-wide association study (GWAS) was performed using a univariate linear mixed model (LMM) to account for relatedness and population structure, as implemented in GEMMA v0.94.1 software (Zhou and Stephens 2012), assessing significance with the Wald test. The statistical model used to test one marker at a time was:

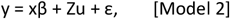

where y is the vector of adjusted phonotypes for all individuals; x is the vector of genotypes at the tested marker; β is the effect of the tested marker; u is a vector of random additive genetic effects distributed according to N(0, Aλτ^-1^), where λ is the ratio of the additive genetic variance and the residual variance τ^-1^, A is the additive relationship matrix, and Z is the incidence matrix (identity matrix in this case); ε is a vector of residuals distributed according to N(0, |τ^-1^), where I is the identity matrix.

GEMMA software implements the Genome-wide Efficient Mixed Model Association algorithm. The first step of the analyses included the estimation of the relatedness matrix. The resulting metrics were included in the second step (GWAS), allowing for the adjustment for both relatedness and population structure. The significance thresholds at the chromosome-wide level (BONF_chr i_) and genome-wide level (BONF_geno_) were obtained using the Bonferroni method that accounts for multiple testing assuming that the number of independent tests was equal to the number of SNPs analyzed (Benjamini and Hochberg 1995). The formulas to obtain thresholds were the following:

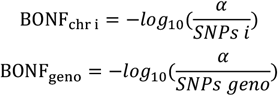

where *SNPs i* is the number of SNPs for chromosome i, and *SNPs geno* is the total number of SNPs at the genome level (i.e., 48,593 autosomal SNPs) and considering *α*=5%. The resulting genome-wide significance threshold was equal to 5.94. GWAS was performed for all of the 26 autosomal chromosomes. The percentage of variance explained by each SNP was calculated as follows:

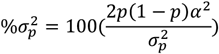

where 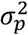 is the phenotypic variance of the trait, *p* is the frequency of the allelic substitution effect of the SNP, and *α* is the estimated allelic substitution effect of the SNP (Sanchez *et al*. 2019). When a given trait was significantly affected by multiple variants, the reported top SNPs are SNPs that have the highest –log 10 (P-value) among the significant SNPs in a 1-Mbp window.

The annotated candidate genes that were closest to the top SNPs were identified using the Ensembl release 104 of the sheep reference genome OAR v3.1 (http://www.ensembl.org).

### LEPR structure and effect of LEPR Genotype

Multiple LEPR protein sequence alignment of several mammals was performed using Weblogo software (Crooks *et al*. 2004) (http://weblogo.threeplusone.com/). Protein sequences are available at NCBI (Mus musculus NP_666258.2, Ratus norvegicus NP_036728.1, Homo Sapiens NP_002294.2, Sus scrofa NP_001019758.1, Bos taurus NP_001012285.2, Ovis aries NP_001009763.1 incomplete sequence) (https://www.ncbi.nlm.nih.gov). The complete LEPR protein sequence in sheep (W5PL31, 1,165 amino acids) was obtained from the UniProt data base (https://www.uniprot.org) from transcript ENSOART00000011314.1 (Ensembl), and we introduced the proline to serine substitution at position 1019.

Analyses of variance were performed to test the effect of the LEPR genotype (i.e., SNP oar3_OAR1_40857869) on BR phenotypes using the MIXED procedure of SAS (version 9.4; SAS Institute Inc., Cary, NC, USA). The dependent variables were adjusted phenotypes for BCS and BCS changes at each physiological stage, as described above. The three possible LEPR genotypes were fitted as a fixed explanatory variable and a significance threshold of p < 0.05 was selected. The Varcomp procedure of SAS was used to fit the genotype effect as random and to estimate the proportion of variance explained by the genotype. The animal was included as a random effect in both the Mixed and Varcomp models.

The effects of the LEPR genotype on BW, BF and BM were independently analyzed at each physiological stage with a linear mixed model using the mixed procedure of SAS. The LEPR genotype was included as a fixed effect, whereas the animal was treated as a random effect. Appropriate following fixed effects, parity, litter size, age at first lambing and year of measurement, were also including in mixed models depending on the trait analyzed.

## Results

### Descriptive statistics

The BCS and BCS changes were significantly affected by the parity and the age at first lambing of the ewe, the litter size, and the year of measurements at several physiological stages (Tables 1 & 2). The first-order interaction parity × litter size was significant for BCS at all physiological stages and was only significant for BCS changes between pregnancy and early suckling (BCS-Pa:L, BCS-Pa:W and BCS-L-Sa. The first-order interaction age at first lambing × litter size was significant for BCS except at lambing, and only significant for BCS changes between pregnancy and weaning (BCS-Pa:W).

**Table 1.** Least-square means for body condition score (BCS) (± standard error) at each physiological stage of ewes according to parity, litter size and age of the ewe at first lambing.

|  |  | %<br>Obs. | BCS-M | BCS-Pa | BCS-Pb | BCS-L | BCS-Sa | BCS-Sb | BCS-W | BCS-Wp |
| --- | --- | --- | --- | --- | --- | --- | --- | --- | --- | --- |
| N Obs. |  |  | 2069 | 2167 | 2167 | 2162 | 1983 | 1317 | 2084 | 1776 |
| Parity | 1 | 47.3 | 2.91 (0.01) a | 2.96 (0.01) a | 2.76 (0.01) a | 2.62 (0.01) a | 2.52 (0.01) a | 2.55 (0.01) a | 2.48 (0.008) a | 2.57 (0.01) a |
|  | 2 | 38.9 | 2.69 (0.01) b | 2.87 (0.01) b | 2.73 (0.01) b | 2.65 (0.01) b | 2.57 (0.01) b | 2.64 (0.02) b | 2.56 (0.01) b | 2.67 (0.01) b |
|  | 3 | 13.8 | 2.73 (0.02) c | 2.95 (0.02) ac | 2.84 (0.02) c | 2.70 (0.02) c | 2.65 (0.02) c | 2.64 (0.02) b | 2.61 (0.01) c | 2.66 (0.02) b |
|  | Sign. |  | *** | *** | *** | *** | *** | *** | *** | *** |
| Litter size | 0 | 13.3 | 2.79 (0.02) ac | 2.93 (0.02) | 2.79 (0.02) ab | 2.75 (0.02) a | 2.82 (0.02) a | 3.04 (0.03) a | 2.97 (0.02) a | 2.89 (0.03) a |
|  | 1 | 20.5 | 2.78 (0.02) ac | 2.94 (0.01) | 2.83 (0.01) b | 2.76 (0.01) a | 2.67 (0.01) b | 2.65 (0.02) b | 2.55 (0.01) b | 2.68 (0.01) b |
|  | 2 | 24.3 | 2.80 (0.01) ab | 2.93 (0.01) | 2.77 (0.01) a | 2.62 (0.01) b | 2.54 (0.01) c | 2.55 (0.02) c | 2.49 (0.01) c | 2.58 (0.02) c |
|  | 3 | 24.6 | 2.78 (0.01) ab | 2.93 (0.01) | 2.76 (0.01) a | 2.63 (0.01) b | 2.47 (0.01) d | 2.42 (0.02) d | 2.39 (0.1) d | 2.52 (0.1) d |
|  | 4 | 17.3 | 2.74 (0.02) c | 2.91 (0.02) | 2.72 (0.02) c | 2.53 (0.01) c | 2.40 (0.01) e | 2.39 (0.02) e | 2.34 (0.01) e | 2.48 (0.02) d |
|  | Sign. |  | * | NS | *** | *** | *** | *** | *** | *** |
| Age at first lambing | 1 | 47.5 | 2.83 (0.01) a | 2.90 (0.01) a | 2.76 (0.01) | 2.66 (0.01) | 2.58 (0.01) | 2.60 (0.02) | 2.53 (0.01) a | 2.56 (0.01) a |
|  | 2 | 52.5 | 2.73 (0.01) b | 2.96 (0.01) b | 2.78 (0.01) | 2.66 (0.01) | 2.57 (0.01) | 2.62 (0.02) | 2.57 (0.01) b | 2.70 (0.01) b |
|  | Sign. |  | *** | *** | NS | NS | NS | NS | *** | *** |
| Year | Sign. |  | *** | *** | *** | *** | *** | *** | *** | *** |
| Parity*Litter size | Sign. |  | *** | * | * | * | *** | *** | *** | *** |
| Age at first lambing*Litter size | Sign. |  | *** | * | *** | NS | * | *** | * | ** |

The significance probabilities were also reported for the year of measurement and first-order interactions. N Obs., number of records for each trait; % Obs., percentage of observations for each class of a given factor. M, Mating; Pa, Early pregnancy; Pb, Two-thirds pregnancy; L, Lambing; Sa, Early suckling; Sb, End of suckling; W, Weaning; Wp, Post-weaning period. Sign., the significance probabilities for each fixed effect of mixed models are provided as : \*\*\* P-value < 0.001, \*\* P-value < 0.01, \* P-value < 0.05; NS, not significant. The lower case letters (a, b, c, d, e) indicate significant differences in the trait between classes of each factor (i.e., values not sharing a common letter are significantly different, as determined by a t-test at $p < 0.05$ ).

**Table 2.** Least-square means for changes in body condition score (BCS) (± standard error) over successive physiological stages of ewes according to parity, litter size and age of the ewe at first lambing.

|  |  | % Obs. | BCS-M:Pa | BCS-Pa:L | BCS-Pa:W | BCS-L:Sa | BCS-W:Wp | BCS-W:M |
| --- | --- | --- | --- | --- | --- | --- | --- | --- |
| N Obs. |  |  | 2060 | 2153 | 2075 | 1978 | 1730 | 1204 |
| Parity | 1 | 47.3 | 0.06 (0.01) a | -0.35 (0.01) a | -0.49 (0.01) a | -0.10 (0.01) a | 0.09 (0.01) a | 0.21 (0.01) a |
|  | 2 | 38.9 | 0.19 (0.01) b | -0.22 (0.01) b | -0.31 (0.01) b | -0.08 (0.01) a | 0.12 (0.01) b | 0.17 (0.02) ab |
|  | 3 | 13.8 | 0.22 (0.02) b | -0.25 (0.02) b | -0.32 (0.02) b | -0.04 (0.02) b | 0.08 (0.02) ab | 0.12 (0.04) b |
|  | Sign. |  | *** | *** | *** | ** | NS | ** |
| Litter size | 0 | 13.3 | 0.12 (0.02) a | -0.18 (0.03) a | 0.07 (0.03) a | 0.09 (0.03) a | -0.02 (0.03) a | -0.12 (0.06) a |
|  | 1 | 20.5 | 0.17 (0.02) ab | -0.17 (0.02) a | -0.38 (0.02) b | -0.10 (0.01) bd | 0.12 (0.02) bc | 0.17 (0.02) b |
|  | 2 | 24.3 | 0.15 (0.02) ab | -0.33 (0.02) b | -0.45 (0.02) c | -0.07 (0.01) b | 0.10 (0.02) b | 0.23 (0.02) c |
|  | 3 | 24.6 | 0.16 (0.02) ab | -0.30 (0.01) b | -0.54 (0.01) d | -0.16 (0.01) c | 0.12 (0.01) bc | 0.27 (0.02) cd |
|  | 4 | 17.3 | 0.18 (0.02) b | -0.37 (0.02) c | -0.57 (0.02) d | -0.13 (0.01) cd | 0.14 (0.02) c | 0.29 (0.02) d |
|  | Sign. |  | NS | *** | *** | *** | *** | *** |
| Age at first lambing | 1 | 47.5 | 0.08 (0.01) a | -0.23 (0.01) a | -0.36 (0.01) | -0.08 (0.01) | 0.04 (0.01) a | 0.12 (0.02) a |
|  | 2 | 52.5 | 0.23 (0.01) b | -0.30 (0.01) b | -0.38 (0.01) | -0.07 (0.01) | 0.15 (0.01) b | 0.22 (0.02) b |
|  | Sign. |  | *** | *** | NS | NS | *** | *** |
| Year | Sign. |  | *** | *** | *** | *** | *** | *** |
| Parity*Litter size | Sign. |  | NS | * | *** | * | NS | NS |
| Age at first lambing*Litter size | Sign. |  | NS | NS | *** | NS | NS | NS |

The significance probabilities were also reported for the year of measurement and first-order interactions. N Obs., number of records for each trait; % Obs., percentage of observations for each class of a given factor. M, Mating; Pa, Early pregnancy; L, Lambing; Sa, Early suckling; W, Weaning; Wp, Post-weaning period; Sign., the significance probabilities for each fixed effect of mixed models are provided as : \*\*\* P-value < 0.001, \*\* P-value < 0.01, \* P-value < 0.05; NS, not significant. The lower case letters (a, b, c, d) indicate significant differences in the trait between classes of each factor (i.e., values not sharing a common letter are significantly different as determined by a t-test at $p < 0.05$ ).

Globally, a significant increase in BCS was observed for BCS-Pb, BCS-L, BCS-Sa, BCS-Sb, BCS-W and BCS-Wp with parity, and a decrease in BCS was observed for BCS-M and BCS-Pa with parity (Table 1). A significant increase in BCS gain was observed for BCS-M:Pa, whereas a decrease in BCS gain was observed for BCS-W:M with the increase in parity (Table 2). A significant decrease in BCS loss was observed for BCS-Pa:L, BCS-Pa:W and BCS-L:Sa with the increase in parity. The litter size effect was significant for BCS-M, BCS-Pb, BCS-L, BCS-Sa, BCS-Sb, BCS-W and BCS-Wp with a decrease in BCS for larger litter size (Table 1). BCS loss increased for BCS-Pa:L, BCS-Pa:W, BCS-L:Sa and BCS gain increased for BCS-W:Wp and BCS-W:M with the increase in litter size (Table 2). The age at first lambing was significant for BCS traits, except between two-thirds pregnancy and end of suckling, with a higher BCS for younger ewes at BCS-M and a lower BCS for younger ewes at BCS-Pa, BCS-W and BCS-Wp (Table 1). A significant increase in BCS gain was observed for BCS-M:Pa, BCS-W:Wp and BCS-W:M, and a significant increase in BCS loss was observed for BCS-Pa:L for ewes that lambed at 2 years of age (Table 2). The year effect was significant (P < 0.01) for all BCS and BCS changes at all physiological stages.

### Genome-Wide Association Studies

The QTLs reaching the chromosome-wide (CW) or genome-wide (GW) thresholds are reported in Table 3. Six QTLs reached the GW significance threshold and 17 reached the CW significance thresholds. Among these 23 QTLs, 16 were related to BCS traits and seven were related to BCS changes. These QTLs were located on 12 different chromosomes. Estimated QTL effects ranged from 1.6 (BCS-W:Wp on OAR25) to 3.9% (BCS-L on OAR1) of the respective phenotypic variance.

**Table 3.** Summary of QTLs detected in GWAS and candidate genes associated with body condition score (BCS) and BCS changes.

| Trait<sup>1</sup> | Chr | Nb SNPs<sup>2</sup> | Top SNP<sup>3</sup> | Position (bp) | -log10(P-value) | SNP effect<sup>4</sup> | Closest gene<sup>5</sup> |
| --- | --- | --- | --- | --- | --- | --- | --- |
| **BCS-Pb** | **1** | **3** | **OAR1_40030112.1** | **38,809,812** | **6.55** | **3.7** | **PGM1** |
| BCS-L | 1 | 1 | OAR1_40030112.1 | 38,809,812 | 5.87 | 3.3 | PGM1 |
| BCS-Pa | 1 | 2 | s30054.1 | 40,524,081 | 5.60 | 2.7 | DNAJC6 |
| **BCS-Sa** | **1** | **3** | **oar3_OAR1_40821987** | **40,821,987** | **6.06** | **2.8** | **LEPR** |
| **BCS-Pb** | **1** | **7** | **oar3_OAR1_40857869** | **40,857,869** | **6.90** | **3.5** | **LEPR** |
| **BCS-L** | **1** | **11** | **oar3_OAR1_40890859** | **40,890,859** | **7.44** | **3.9** | **LEPR** |
| **BCS-Pb** | **1** | **2** | **OAR1_42228129.1** | **42,228,129** | **6.03** | **2.9** | **MIER1** |
| BCS-Sb | 2 | 1 | OAR2_204233702.1 | 192,716,355 | 5.51 | 2.5 | MYO1B, STAT1 |
| BCS-Pa:W | 3 | 1 | OAR3_115100887.1 | 115,100,887 | 5.24 | 2.4 | SYT1 |
| BCS-Pb | 8 | 1 | s48527.1 | 13,120,875 | 4.84 | 2.0 | TPD52L1 |
| BCS-Pa | 10 | 1 | OAR10_9706403.1 | 11,281,919 | 5.00 | 2.0 | PCDH8 |
| BCS-Pa | 15 | 1 | OAR15_26097184.1 | 24,911,875 | 4.82 | 2.3 | NXPE4 |
| BCS-Sa | 15 | 1 | OAR15_87285774.1 | 78,544,042 | 5.22 | 2.3 | OR10W1 |
| BCS-Sa | 15 | 1 | OAR15_87912118.1 | 87,912,118 | 4.65 | 2.2 | - |
| **BCS-W:Wp** | **16** | **2** | **OAR16_33763548.1** | **31,368,878** | **6.59** | **2.6** | **CCL28** |
| BCS-W:Wp | 16 | 1 | OAR16_34857607.1 | 34,857,607 | 4.99 | 1.9 | DAB2 |
| BCS-Pa:L | 16 | 1 | OAR16_46544413.1 | 42,819,279 | 5.03 | 2.0 | CDH6 |
| BCS-Sa | 17 | 1 | s70069.1 | 14,052,727 | 4.47 | 2.3 | FREM3, GAB1 |
| BCS-L | 18 | 1 | OAR18_31578626.1 | 30,304,217 | 4.65 | 2.4 | NRG4 |
| BCS-M:Pa | 22 | 1 | OAR22_24239807.1 | 24,239,807 | 4.39 | 1.9 | SORCS3 |
| BCS-Pa:L | 24 | 1 | OAR24_22245800.1 | 20,494,588 | 4.20 | 1.9 | HS3ST2 |
| BCS-W:Wp | 25 | 1 | s68395.1 | 32,693,219 | 4.43 | 1.6 | KCNMA1 |
| BCS-Pa | 25 | 1 | OAR25_39067458.1 | 37,133,591 | 4.75 | 2.5 | - |

<sup>1</sup>M, Mating; Pa, Early pregnancy; Pb, Two-thirds pregnancy; L, Lambing; Sa, Early suckling; Sb, End of suckling; W, Weaning; Wp, Post-weaning period. <sup>2</sup>Number of significant SNPs in the 1-Mb window. <sup>3</sup>The reported top SNPs are SNPs that have the highest -log10(P-value) among the significant SNPs that are in 1-Mb windows. SNPs for which -log10(P-value) reached the genome-wide significance threshold (> 5.98) are reported in bold; the other reported SNPs reached the chromosome-wide significance threshold. <sup>4</sup>Percentage of variance explained by SNP. <sup>5</sup>Annotated protein coding genes closest to the top SNP of the QTL region. Chr, chromosome.

Five of the six significant associations reached the GW significance threshold mapped on OAR1 and were associated with BCS traits (Table 3, Figure 1). Two additional QTLs reached the CW threshold localized on OAR1. All the significant SNPs mapped on OAR1 were located between 38.80 and 42.22 Mb. A first QTL region mapped on OAR1 at 38.80Mb was associated with BCS-Pb and BCS-L. A second QTL region on OAR1 located between 40.26 and 41.56 was associated with four BCS traits (BCS-Pa, BCS-Pb, BCS-L and BCS-Sa). A third QTL region located on OAR1 at 42.22 Mb was associated with BCS-Pb. Details of the second QTL region located on OAR1 are given in Table 4. This QTL region contained 14 SNPs, significantly associated with at least one BCS trait. Among these 14 SNPs, four SNPs mapped in the LEPR gene, including three SNPs localized in the intronic sequence and one SNP in the coding sequence.

**Figure 1.**
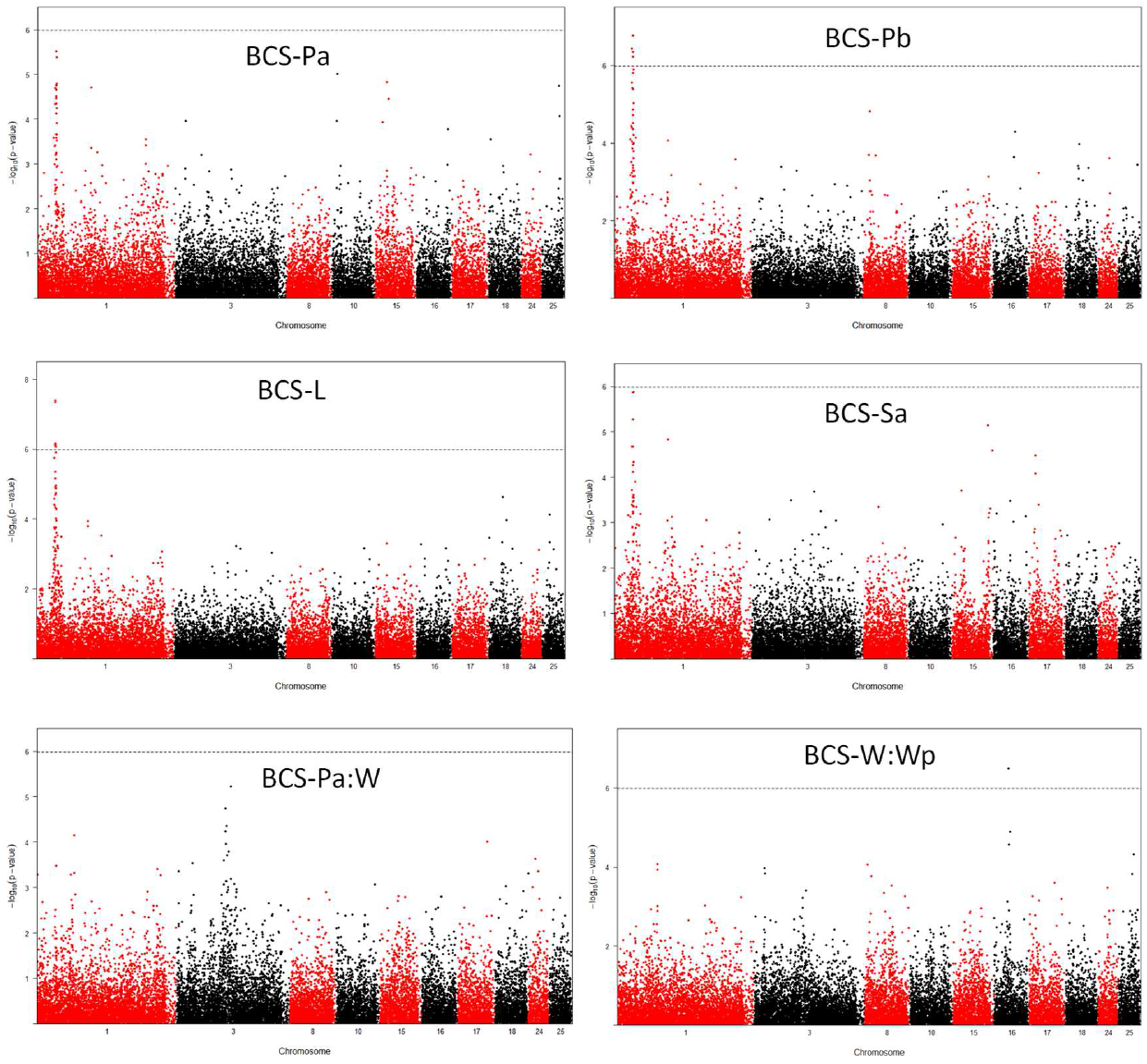
Chromosome plots of body condition score (BCS) and BCS changes. The –log_10_(p-value) for all SNPs were plotted for chromosomes 1, 3, 8, 10, 15, 16, 17, 18, 24 and 25. The dashed line indicates the genome-wide significance threshold (BONF_gen_ = 5.94); The chromosome-wide significance thresholds were OAR1: 5.02, OAR3: 4.96, OAR8: 4.57, OAR10: 4.52, OAR15: 4.49, OAR16: 4.45, OAR17: 4.42, OAR18: 4.43, OAR24: 4.14, OAR25: 4.26. Pa, Early pregnancy; Pb, Two-thirds pregnancy; L, Lambing; Sa, Early suckling; W, Weaning; Wp, post-weaning.

**Table 4.** Details of the 14 single nucleotide polymorphisms (SNPs) detected in the common OAR1 region associated with four BCS traits.

| SNP | Position | MAF | P-value |  |  |  | Gene | Variant type |
| --- | --- | --- | --- | --- | --- | --- | --- | --- |
|  |  |  | BCS-Pa | BCS-Pb | BCS-L | BCS-Sa |  |  |
| OAR1_41661218.1 | 40,268,642 | 0.39 | *3.05E-05* | *1.91E-05* | **7.40E-07** | *5.71E-04* | no | intergenic |
| s30054.1 | 40,524,081 | 0.09 | *2.51E-06* | *3.28E-04* | *2.75E-03* | *3.21E-04* | no | intergenic |
| OAR1_42038601.1 | 40,539,479 | 0.36 | *2.72E-05* | *3.06E-05* | *4.58E-06* | *1.41E-02* | no | intergenic |
| oar3_OAR1_40821987 | 40,821,987 | 0.35 | *3.42E-05* | *2.74E-06* | *1.88E-05* | **8.63E-07** | LEPR | intron |
| oar3_OAR1_40828247 | 40,828,247 | 0.44 | *1.55E-05* | **3.61E-07** | **6.81E-07** | *3.50E-06* | LEPR | intron |
| oar3_OAR1_40848298 | 40,848,298 | 0.32 | *2.72E-05* | **1.27E-07** | **4.11E-08** | *4.32E-05* | LEPR | intron |
| oar3_OAR1_40857869 | 40,857,869 | 0.32 | *2.72E-05* | **1.27E-07** | **4.11E-08** | *4.32E-05* | LEPR | missence |
| oar3_OAR1_40865586 | 40,865,586 | 0.32 | *2.72E-05* | **1.27E-07** | **4.11E-08** | *4.32E-05* | no | intergenic |
| oar3_OAR1_40872161 | 40,872,161 | 0.50 | *1.10E-02* | *4.60E-04* | *6.76E-06* | *3.83E-04* | no | intergenic |
| oar3_OAR1_40890859 | 40,890,859 | 0.32 | *6.15E-05* | **3.31E-07** | **3.61E-08** | *6.09E-05* | no | intergenic |
| oar3_OAR1_41137040 | 41,137,040 | 0.31 | *1.33E-05* | *1.17E-06* | **6.10E-07** | *2.50E-04* | no | intergenic |
| OAR1_43022391.1 | 41,454,757 | 0.28 | *2.61E-04* | *8.85E-05* | *1.04E-06* | *2.55E-04* | no | intergenic |
| OAR1_43083702.1 | 41,512,082 | 0.62 | *1.05E-03* | *1.41E-04* | *1.02E-05* | *1.15E-06* | no | intergenic |
| oar3_OAR1_41564208 | 41,564,208 | 0.61 | *3.34E-06* | *8.10E-05* | **7.10E-07** | *3.77E-04* | no | intergenic |

SNP, Name of the single nucleotide polymorphism; MAF, minor allele frequency; P-value, the p-values reported correspond to the unadjusted p-value from the Wald test. In bold, SNP significant at the genome-wide threshold; in italics, non-significant SNPs at the chromosome- wide threshold. Gene, gene is reported when SNP overlapped with a gene. LEPR, Leptin receptor.

One additional significant association was also detected with BCS-Pb and BCS-L and mapped on OAR8 and OAR18, respectively (Table 3). Only one QTL was associated with BCS-Sb and localized on OAR2. Three additional QTLs were associated with BCS-Pa and mapped on OAR10, OAR15 and OAR25 (Table 3). Similarly, three additional QTLs were associated with BCS-Sa and mapped on OAR15 at 78.54 and 87.91Mb and OAR17. Three QTLs were significantly associated with BCS-W:Wp and mapped on OAR16 at 31.36 and 34.87Mb and on OAR25. The QTL associated with BCS-W:Wp located on OAR16 at 31.36Mb reached the GW threshold (Table 3, Figure 1). Two QTLs were associated with BCS-Pa:L and localized on OAR16 at 42.81Mb and OAR24. Finally, one QTL was associated with BCS-M:Pa and mapped on OAR22, and one QTL was associated with BCS-Pa:W and mapped on OAR3.

### Leptin receptor structure

Among SNPs significantly associated with BCS traits and located in the LEPR gene, only a single SNP (i.e., oar3_OAR1_40857869) mapped to the coding region of the gene (Figure 2A). The mutation induced a non-synonymous change in amino acid. Modification of the C base in the reference sequence (OAR1, 40857869 bp) to a T encoded a proline to serine substitution at position 1019 (p.P1019S) of the leptin receptor protein (Figure 2B). The mutation is located within the cytoplasmic domain and replacement of proline by serine causes a polar to non-polar amino acid substitution. While tyrosine phosphorylation sites of the cytoplasmic domain were highly conserved across species (i.e., Y986, Y1078, Y1141), this sequence of the LEPR protein was poorly conserved across species in the region surrounding the mutation (Figure 2C).

**Figure 2.**
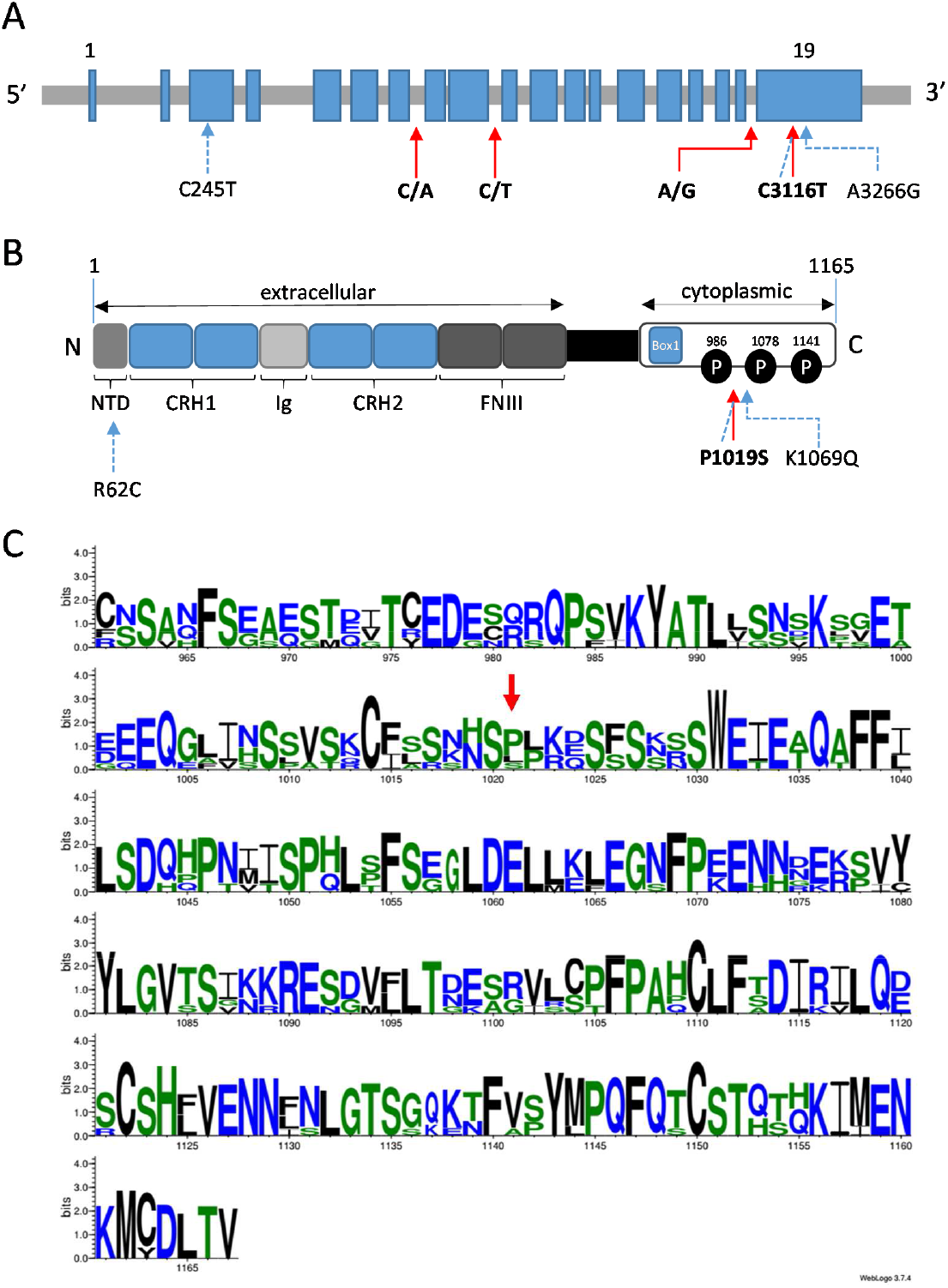
The leptin receptor (LEPR) gene and protein structure and polymorphisms in sheep. (A) Schematic nucleotide sequence structure of the LEPR gene and sheep polymorphisms previously reported to be associated with reproductive phenotypes (dashed arrows) or identified in the present study (solid arrows). Arrows indicate the position within the gene and the type of base pair exchange. Numbers, when reported, indicate position within the cDNA sequence ENSOART00000011314.1. (B) LEPR protein structure based on the long form (1,165 amino acids) and polymorphisms previously reported (dashed arrows) or identified in the present study (solid arrow). Arrows indicate the position within the protein and the type of amino acid exchange. NTD: N-terminal domain; CRH: cytokine receptor homology; Ig: immunoglobulin-like domain; FNIII: fibronectin type III; Leptin binds to its homodimer receptor through CRH2 and activates downstream effectors through Box1 domain and phosphorylation (P) of Y-residues (Y986, Y1078, Y1141) (adapted from Berger et al. (BERGER AND KLÖTING 2021)). (C) Multiple alignment of the LEPR protein sequences from mouse (NP_666258.2), rat (NP_036728.1), human (NP_002294.2), pig (NP_001019758.1), cattle (NP_001012285.2) and sheep (W5PL31) species with Weblogo software (Crooks *et al*. 2004). Only the C-terminal end of the protein is represented. A red arrow indicates the P1019 position in sheep. Numbering on the X-axis resulted from the multiple alignment and not the ovine sequence.

### Effects of the LEPR genotype

The frequencies of wild-type (C/C), heterozygous (C/T) and homozygous (T/T) carriers were 47%, 43% and 10%, respectively. Analysis of variance confirmed that the T mutation involved a significant increase in BCS at all physiological stages of the productive cycle, as shown by the average BCS values in each genotype (Figure 3). Wild-type (C/C) ewes showed the lowest BCS all along the productive cycle, whereas heterozygous (C/T) ewes showed intermediary BCS levels and homozygous (T/T) ewes exhibited the highest BCS. In addition to BCS, the LEPR genotype had significant effects on the body weight, back fat and muscle thickness depending of the physiological stages (Table 5). The homozygous (T/T) ewes had significantly higher body weight than wild-type (C/C) ewes from mating to lambing. The homozygous (T/T) ewes had significantly higher back fat thickness than wild-type (C/C) ewes at mating and end of pregnancy. The homozygous (T/T) ewes had significantly higher back fat muscle than wild-type (C/C) ewes at the last third of pregnancy.

**Figure 3.**
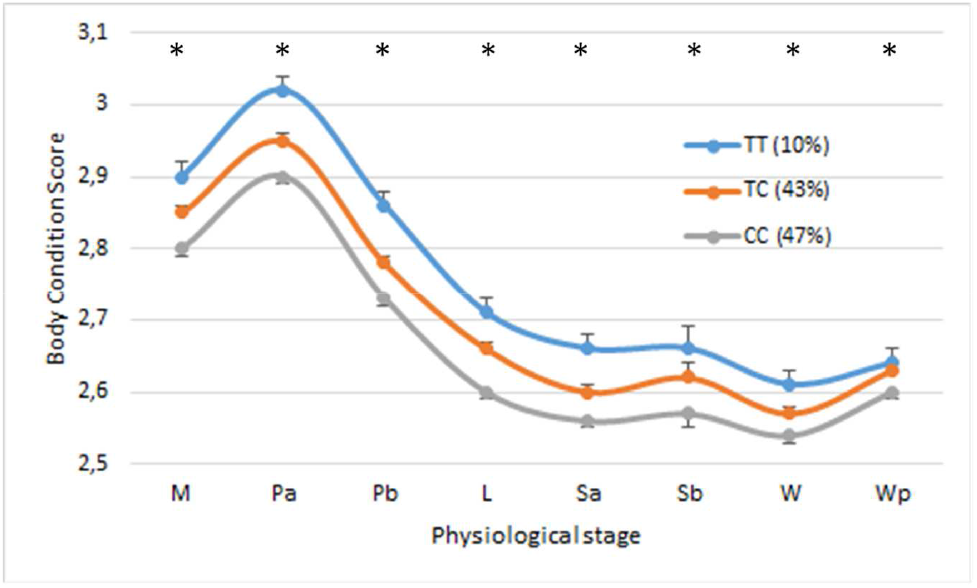
Effect of LEPR genotype on body condition score throughout the productive cycle of ewes. Values represent the lsmeans for body condition score and were obtained from the mixed model with repeated measurements over three successive productive cycles. The asterisk shows the significant overall effect of the LEPR genotype on the trait at p < 0.05. The percentage of ewes for each genotype is given between parentheses. M, Mating; Pa, Early pregnancy; Pb, Two-thirds pregnancy; L, Lambing; Sa, Early suckling; Sb, Mid suckling; W, Weaning; Wp, post-weaning.

**Table 5.** Effect of the LEPR genotype on body weight (BW), back fat depth (BF) and back muscle depth (BM) of ewes throughout the productive cycle.

|  | TT | TC | CC | Sign. |
| --- | --- | --- | --- | --- |
| n (%) |  |  |  |  |
| BW-M | 53.20 (0.47) a | 52.4 (0.26) ab | 51.9 (0.25) b | * |
| BW-Pa | 56.50 (0.5) a | 55.4 (0.28) ab | 54.86 (0.27) b | * |
| BW-Pb | 61.10 (0.52) a | 60.4 (0.29) a | 59.6 (0.28) b | * |
| BW-L | 59.5 (0.75) a | 58.3 (0.57) ab | 57.8 (0.56) b | * |
| BW-Sa | 58.6 (0.58) | 58.2 (0.34) | 57.8 (0.33) | NS |
| BW-Sb | 60.2 (0.73) | 59.8 (0.45) | 59.6 (0.44) | NS |
| BW-W | 56.2 (0.55) | 56.0 (0.30) | 55.7 (0.29) | NS |
| BW-Wp | 57.5 (0.54) | 57.3 (0.31) | 56.8 (0.30) | NS |
| BF-M | 5.44 (0.3) a | 5.1 (0.2) ab | 4.9 (0.2) b | * |
| BF-Pb | 5.0 (0.2) a | 4.3 (0.2) b | 4.2 (0.2) b | ** |
| BF-W | 4.5 (0.3) | 4.3 (0.2) | 4.3 (0.2) | NS |
| BM-M | 22.5 (0.7) | 22.3 (0.5) | 22.3 (0.5) | NS |
| BM-Pb | 21.5 (0.7) a | 19.7 (0.5) b | 19.8 (0.5) b | * |
| BM-W | 19.6 (0.9) | 19.4 (0.7) | 18.9 (0.7) | NS |

Values reported are lsmeans ( ± standard error) obtained from the mixed model with repeated measurements over successive productive cycles. Sign. indicates the significant overall effect of the LEPR genotype on the trait at \* $p < 0.05$ , \*\* $p < 0.01$ . NS, not significant. Different letter superscripts (a, b, c) show a significant difference between genotypes at $p < 0.05$ . n (%), proportion of ewes in each genotype; M, Mating; Pa, Early pregnancy; Pb, Two-thirds pregnancy; L, Lambing; Sa, Early suckling; Sb, End of suckling; W, Weaning; Wp, Post-weaning period. BW, body weight in kg; BF, back fat thickness in mm; BM, back muscle thickness in mm.

## Discussion

Considering the existing genetic variability for energetic body reserve traits previously reported in sheep (Banos *et al*. 2005; Walkom *et al*. 2014a; Macé *et al*. 2018), the aim of the present study was to provide the first characterization of the genetic architecture that controls body condition in productive ewes. This was achieved by a genome-wide association study (GWAS) of a set of body reserve (BR) traits measured at key physiological stages of the productive cycle in Romane ewes.

### Body reserve dynamics

Body reserve changes throughout ewe productive cycles found for the sheep population used in the present study have been described and discussed in detail by Macé et al. (Macé *et al*. 2018; Macé *et al*. 2019). Sources of variation affecting BR and found in the present study were fully consistent with those previously described in our experimental conditions, even if the number of sheep was slightly different since we kept only phenotyped and genotyped ewes. Briefly, body reserve changes over time were highly influenced by physiological stages. Generally, BR accretion was observed from weaning to early pregnancy, whereas BR mobilization was observed from two-thirds pregnancy to weaning, which is probably linked to the negative energy balance induced by the increase in energetic requirements during pregnancy and suckling periods. The BR levels and BR changes over time reported in the present study were also affected by the biological effects of parity, litter size and age of ewes at first lambing. The increase in BR mobilization and accretion with litter size were consistent with the higher energy requirements induced by multiple litters and the high genetic correlation between BR mobilization and accretion processes previously reported by Macé et al. (Macé *et al*. 2018). Interestingly, the increase in BR accretion and the decrease in BR mobilization with parity suggested that ewes, thanks to their metabolic experience in the previous cycles, may develop strategies either linked with BR management or feeding to limit negative effects of excessive BR mobilization.

### QTLs for BR traits

As far as we know, this is the first study on ruminants that maps QTLs for BR levels at several key physiological stages and BR changes over time in ewes. Indeed, many QTLs have been previously found for conformation traits and carcass fatness in sheep (Walling *et al*. 2004; Matika *et al*. 2016; Garza Hernandez *et al*. 2018; Hu *et al*. 2018), but not yet for BR traits in live productive ewes. In the present study, we were not only interested in BR levels assessed through body condition score (BCS) but also in body reserve dynamics (BRD) assessed through BCS changes over time (i.e., between physiological stages). The GWAS analyses resulted in the mapping of many QTLs associated with BR levels (OAR1, 2, 8, 10, 15, 17, 18, 25) and BR changes (OAR3, 16, 22, 24, 25).

The findings were of particular interest for associations mapped on chromosome 1 (OAR1). The QTL regions detected on OAR1 were both associated with several correlated traits and showed a high level of significance. These QTL regions on OAR1 were associated with body reserve levels at pregnancy, lambing and early suckling. Interestingly, we previously reported that BR mobilization occurred between two-thirds pregnancy and weaning in ewes reared in the extensive conditions of the La Fage farm (Macé *et al*. 2019). Thus, the QTLs on OAR1 may be associated with BR levels during the BR mobilization process. Overlapping on OAR1 for QTLs associated with BR levels at several physiological stages of the mobilization period was consistent with the high genetic correlations previously reported for BR levels between physiological stages (Macé *et al*. 2018).

The common QTL region on chromosome 1 (40.26 to 41.56 Mb) associated with BR levels at four key physiological stages, harbored several highly significant SNPs, including four SNPs overlapped with the most interesting candidate gene: *LEPR* (leptin receptor). The *LEPR* gene codes for the receptor of the leptin hormone. The leptin hormone, secreted by adipose tissue, and the leptin receptor have been widely described for their major role in energy regulation (Speakman *et al*. 2008; Berger and Klöting 2021; Bouchard 2021). Mutations in leptin and LEPR genes have been reported to cause obesity in human and animal models (Chagnon *et al*. 1999; Yiannakouris *et al*. 2001; Israel and Chua 2010; Ghalandari *et al*. 2015; Berger and Klöting 2021). In sheep, by using a candidate gene approach, Haldar et al. (Haldar *et al*. 2014) described three mutations in LEPR associated first with reproductive traits. One of these mutations found in the Davisdale sheep breed is located at 40,857,869 bp on chromosome 1 and corresponds to the SNP oar3_OAR1_40857869 significantly associated with BR levels in the present study. This mutation in the coding region of the gene, modifying a cytosine to a thymidine, causes an amino acid change in the cytoplasmic domain, implying a substitution from proline to serine (p.P1019S), which is not observed in other species. This variation causes a polar to a non-polar amino acid substitution and may alter a potential phosphorylation site from the serine directly preceding it, as suggested by Haldar et al. (Haldar *et al*. 2014). Whether this would alter the receptor function remains unknown because the potential role of phosphorylation of serine at amino acid 1018, which is highly conserved across species, has not been examined to date. We also cannot exclude the possibility that additional polymorphisms may exist in the LEPR gene in the Romane sheep breed, which could alter LEPR functional activity.

Many studies have documented the essential role of the LEPR protein in energy regulation. Various mutations in the LEPR gene leading to leptin signaling deficiency through disruption in the LEPR function resulted in many cases in obesity/diabetes phenotypes in humans and rodents (Chagnon *et al*. 1999; Yiannakouris *et al*. 2001; Israel and Chua 2010; Ghalandari *et al*. 2015; Berger and Klöting 2021). In pigs, a missense mutation in the *LEPR* gene was also associated with higher fatness levels (Uemoto *et al*. 2012; Ros-Freixedes *et al*. 2016) and antagonistic maternal and direct effects on body weight (Sole *et al*. 2021). In sheep, in addition to the effect on reproductive traits, Haldar et al. (Haldar *et al*. 2014) also found an effect of the LEPR p.P1019S mutation on adult body weight, with ewes with the mutation being heavier than the other ewes. They also reported that for ewes homozygous for the LEPR p.P1019S mutation, BCS at 18 months of age was 5 to 10% greater than for the wild-type ewes. This is in accordance with our present results showing higher body weight and fatness levels in ewes carrying the LEPR p.P1019S mutation. When using BCS as proxy of fatness, higher fatness levels in ewes carrying the LEPR p.P1019S mutation was observed all along the productive cycle and not only at the key physiological stages of BR mobilization. The range of differences in BCS between genotypes was in agreement with the individual variability that we previously reported in our farming conditions (Macé *et al*. 2019). There are multiple underlying causes of obesity phenotypes in LEPR mutants, including hyperphagia, increased lipogenesis, or increased feed efficiency (Israel and Chua 2010). Considering the effects of the LEPR p.P1019S mutation observed in the present study, it may be hypothesized that the present mutation may result in a leptin signaling deficiency in sheep and increased lipogenesis.

Two additional QTL regions were found on OAR1, flanking the QTL region containing the LEPR gene described above, and both are associated with BR levels during pregnancy. Interestingly, one of these QTL region maps close to the gene encoding PGM1 (phosphoglucomutase 1). The protein *PGM1* is known to have a central role in gluconeogenesis and glycolysis in humans (Putt *et al*. 1993). Involvement of PGM1 in energetic metabolism and the QTL found close to this gene in the present study makes PGM1 an additional potential candidate gene for BR regulation in sheep.

Nine additional QTL regions were associated with BR levels but only a single SNP reached the chromosome-wide significance level for each of these regions. These QTLs were mainly associated with BR levels during the BR mobilization period, and only QTLs associated with BR at early pregnancy were found for the BR accretion period. Indeed, no QTLs were found for BR levels post-weaning and at mating. Concerning BR changes over time, only a few QTLs were found. The low number of QTLs associated with BR changes over time could be due to a lower genetic variability for these traits compared to BR levels showing higher genetic variability (Macé *et al*. 2018). Among the four QTLs associated with BR gain, only one QTL region reached the genome-wide significance threshold (OAR 16). Several coding genes are located close to the fine location of the QTL in these regions, although no scientific evidence has yet suggested their involvement in regulating energy balance and/or body fatness. Among these genes, the gene DAB2 codes for a protein that acts as a regulator of the activity of protein serine/threonine kinase and could be involved in the leptin signaling pathway.

## Conclusions

The work reported here is the first SNP-based QTL detection for body reserve traits at key physiological stages in productive ewes. We reported various QTLs, including a major QTL on OAR1 associated with BR levels during the BR mobilization period. This QTL region on OAR1 harbors an interesting candidate gene, LEPR, previously described as being associated with obesity and energy regulation in several species. The present identification of a candidate mutation in the LEPR gene provides new opportunities for a deeper understanding of the genetic regulation involved in body reserve management in mammals. Further studies will be developed to investigate functional consequences on the LEPR protein of the identified mutation and to search for potential additional genetic variants in this gene. The impact of this mutation on production traits will also be investigated before considering this genetic variant in small ruminant breeding schemes in order to improve adaptation.

## Data Availability Statement

The phenotypes and genotypes data used for this study are available in the Zenodo public repository (Hazard *et al*. 2021) (DOI: 10.5281/zenodo.5729197).

## Acknowledgments

The authors are indebted to the entire staff of the INRAE La Fage Experimental Farm for their active role in collecting experimental data.

